# Electrical Stunning does not affect Brain Tissue Integrity and Stress Response in Larval Zebrafish

**DOI:** 10.64898/2026.09.04.749313

**Authors:** David-Samuel Burkhardt, Kevin Schultz, Michael Glaser, Stefanie Krais, Demis Maile, Heinz-R. Köhler, Rita Triebskorn, Aristides B. Arrenberg

## Abstract

Electrical stunning is a humane euthanasia method for larval zebrafish, and a suitable euthanasia protocol has recently been established. Here, we assess the associated stress response and tissue integrity. When compared to MS-222 overdose and hypothermia, electrical stunning caused no significant Hsp70 increase. Brain tissue was preserved and muscle tissue affected only slightly. Our results support the suitability of electrical stunning for humane euthanasia and artifact-free post-mortem analyses.

## Main

Electrical stunning is one of the four legal methods for euthanizing zebrafish within the EU. Although it appears to be a promising and humane alternative to currently used methods, such as anesthetic overdose and rapid cooling, it is not yet widely established as a standardized method. This lack of adoption is – in part - due to insufficient scientific data regarding the physiological and neurophysiological effects of electrical stunning and the absence of commercially available, adequate equipment (Köhler et al., 2017; Lidster et al., 2017). So far, only a few studies investigated effects of electrical stunning on zebrafish and its application as a method of euthanasia (Burkhardt et al., 2025; Mocho et al., 2022; Saarinen et al., 2025). We recently published a study investigating both the behavior and neuronal activity during electrical stunning in 4-day-post-fertilization (dpf) zebrafish larvae. Our findings provided evidence that this method is more rapid in its onset (<1s till loss of consciousness), more effective and, therefore, more humane compared to an anesthetic overdose or rapid cooling, when using the proposed parameter set (Burkhardt et al., 2025). However, two questions remained unanswered so far: First, although electrical stunning has an immediate onset, causing fish to lose equilibrium, behavior and coordinated neuronal activity within a second, the extent of stress larvae experience during exposure is unclear. Second, it is unknown how much the method damages the larval tissues, which could potentially complicate or even prevent subsequent postmortem experiments such as toxicological and morphological studies, or transcriptomics (Driessen et al., 2013; Huang et al., 2020).

Here, we investigated both the stress response after electrical stunning, measured by stress protein levels (hsp70), which in the past has been successfully used as an biochemical indicator of early stress responses to noxious stimuli (Bai et al., 2022; Scheil et al., 2010; Xiong et al., 2017), as well as potential tissue alterations and damages using whole-larvae paraffin sections stained with hematoxylin-eosin. To put the results into a broader context and to compare stress responses after the application of different euthanasia methods, hsp70 measurements were also performed in larvae treated with a high dose of MS-222 (1000mg/l), hypothermia (0-2°C), a heat shock (39°C) as well as in untreated larvae. This research aims to further validate electrical stunning as an alternative to other established methods of euthanasia.

To examine the effect of electrical stunning on different zebrafish tissues, 4 dpf larvae were fixed post-treatment with 2 % glutaraldehyde (in 0.1 M cacodylate buffer, pH 7.4). After fixation, samples were dehydrated with ethanol and routinely processed for embedding in technovit resin. To assess alterations and potential neuronal damage due to electrical stunning, we analyzed the midbrain of the larvae for histopathological alterations. Tests were conducted in both fish lines (*mitfa*^*w2/w2*^ and *Tg(elavl3:H2B-GCaMP6f)jf7*) used in our recent electrical stunning study (Burkhardt et al., 2025). Hematoxylin and eosin staining of brain sections of both treated and control larvae generally displayed a well-preserved architecture of the brain (Figure1, Panels iA-B & iiA-B). Major brain regions, including the forebrain, midbrain, and hindbrain, were clearly distinguishable and maintained their normal anatomical structure. Neuropil regions showed no alterations or significant differences in any group, i.e. intercellular space appeared similar in size and coloration did not change (see regions 2&8 in Figure1). Both cell diameter and cell membrane integrity were apparently not affected by the treatment (see region 8 in Figure 1, Panels iiA-B). Furthermore, no signs of karyolytic processes were detected, indicating that the cells were still structurally intact (see region 8 in Figure 1). None of the animals had developed visible edema. Further, skeletal musculature was investigated, as these muscles are known to be activated by the AC field (Burkhardt et al., 2025), to identify potential damage or morphological changes. We specifically focused on the integrity of myomeres (see structure III in Figure 1) and myosepta (see arrow 3 in Figure 1), intercellular spaces, and the presence of hemorrhages. In the control group, individual myosepta were clearly visible and maintained a normal trapezoidal shape, with individual muscle fibres forming myomeres (see structure III in Panels iC-D in Figure 1). Myomeres and muscle fibres lay adjacent to each other without large intercellular spaces. In contrast, treated groups showed deformed myosepta, with the characteristic trapezoidal shape sometimes distorted, and larger and more numerous intercellular spaces, primarily located proximally (see structure III in Panels iiC-D in Figure 1). These changes were particularly visible in the mitfa^w2/w2^ fish experiment (see Panel 2C in Figure 1). However, no muscle fiber ruptures were observed in any group. No hemorrhages or signs of karyolysis or hypertrophy were found. Investigation of the skin in control larvae revealed both the superficial and basal stratum to be clearly visible and intact, exhibiting a slightly dome-shaped structure without any signs of damage. Conversely, almost all larvae in the treated groups displayed peridermal desquamation and cell detachments within the superficial stratum, alongside signs of hypertrophy. The basal stratum, however, consistently appeared unaffected and remained intact.

**Figure 1.**
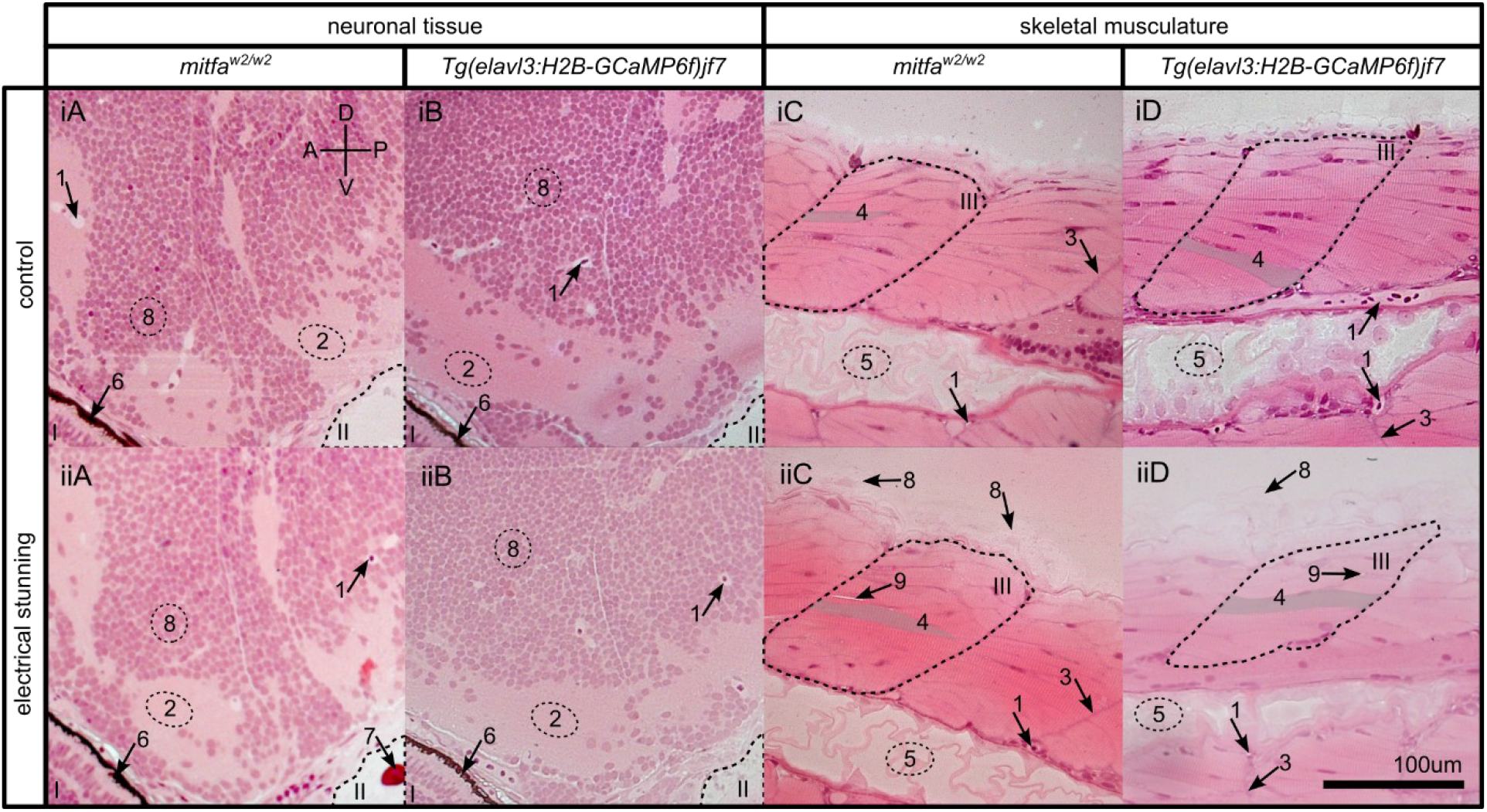
H&E staining of zebrafish neuronal and skeletal tissue for electrically stunned and untreated larvae (sagittal view). Panels 1A-B & 2A-B: forebrain and midbrain. Panels 1C-D & 2C-D: tail musculature. Control and treated groups showed no visible alterations or damage in neuronal tissue. (see regions 2 & 8 in Panels 1A-B & 2A-B). Skeletal musculature showed deformed myosepta and myomeres (see structure III & arrow 3 in Panels 2D-C) in treated group. I: eye; II: inner ear; III: myomere; 1: blood vessel with erythrocyte; 2: axons and dendrites (neuropil); 3: myosepta; 4: myofibrils (shaded area); 5: notochord; 6: retinal pigmented epithelium; 7: rostral otolith; 8: nuclei of neurons.

Stress protein analysis (hsp70) revealed no significant increase in hsp70 levels in larvae following electrical stunning compared to untreated controls. We also investigated hsp70 levels in zebrafish larvae subjected to hypothermic shock (0 - 2 °C for 1h) and an overdose of the anesthetic MS-222 (1000mg/l for 1h) to compare stress responses across different common euthanasia methods (Figure 2A). No significant increases were observed relative to the control group, although the hypothermic shock group exhibited the highest hsp70 levels, followed by MS-222-treated larvae, and the electrically stunned group showed the lowest levels (Figure 2B). To confirm our hsp70 analysis method, a group of larvae was treated with hyperthermic shock (39 °C for 1h), resulting in a significant increase in hsp70 levels compared to controls (p = 0.029, Mann-Whitney-Wilcoxon’s U-test). We further examined whether hsp70 levels in electrically stunned larvae would rise post-stunning - despite larvae being considered unconscious and dead after the first second of electrical exposure. Indeed, one hour after euthanasia via electrical stunning, hsp70 levels were almost twice as high as in larvae analyzed immediately after treatment. This strongly indicates that, although larvae are considered dead based on prior behavioral and neurophysiological measurements (Burkhardt et al., 2025), their cells remain sufficiently intact to induce gene transcription and translation, and thus produce elevated hsp70 levels – at least within 1 hour after treatment.

**Figure 2.**
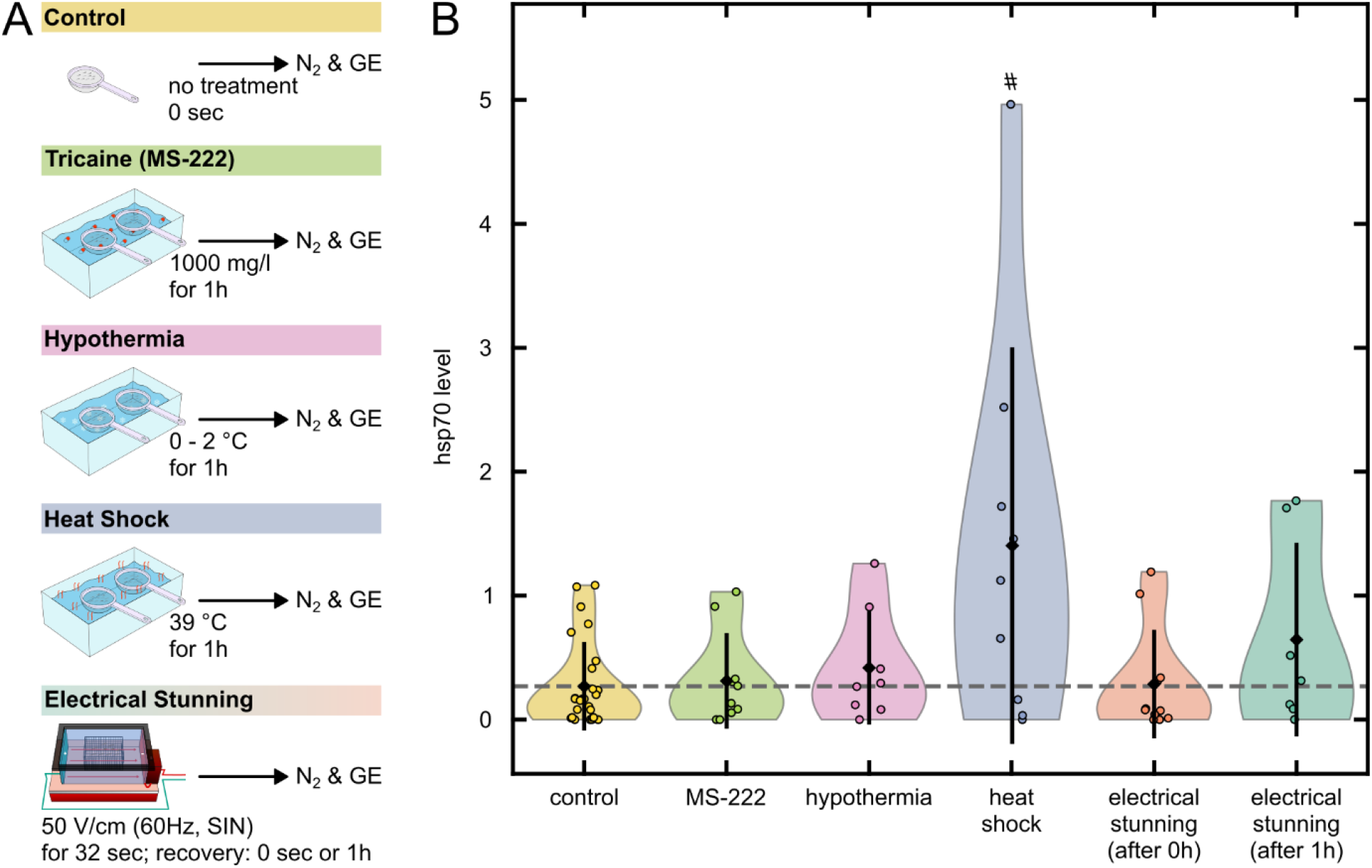
hsp70 measurements after different treatments. A) Protocols for different treatments. N_2_ = liquid nitrogen, GE = gel electrophoresis. B) hsp70 levels (mean ± SD) in larval zebrafish after different treatments. Gray dashed line indicates hsp70 level of control group; # indicates significance at p < 0.05 (Mann-Whitney-Wilcoxon’s U-test).

In Burkhardt et al. (2025), we speculated that death caused by electrical stunning might occur due to ionic imbalance, with the electrical field initially rupturing cell membranes. However, this study cannot confirm cell damage - in fact, neuronal tissue appears largely unaffected (cell size is similar and nuclei are intact) and muscular tissue was only mildly affected by electrical stunning. While this is good news for the electrical stunning method itself, allowing post-mortem experiments such as toxicological, morphological, and potentially even transcriptomics studies, it does not provide any direct hint to the mechanism underlying electrically induced organismal death. The histological results also match our previous observations that AC is the least harmful and best-suited voltage type for euthanizing larval zebrafish (Burkhardt et al., 2025). However, it must be noted that this holds true only when analyzing tissue directly after electrical stunning. Due to the previously observed slow-propagating calcium waves following electrical stunning and progressive rigor mortis, the tissue will further undergo necrosis over time, making artifact-free morphological analyses more difficult. Our hsp70 analysis draws a more distinct picture: electrical stunning induces no measurable increase in cellular stress directly after the treatment, but hsp70 levels rise afterwards. This timeline suggests a temporally distinct organismal death, as judged by lacking behavioral or neurophysiological signs of life at the organismic level, followed by delayed cellular stress responses in cells “surviving” organismal death for minutes to hours (Burkhardt et al., 2025). Although differences in hsp70 levels across different established methods of euthanasia were not statistically significant, the observed low hsp70 levels and mostly preserved tissue integrity confirms electrical stunning as a humane and rapid alternative to hypothermic shock and MS-222 overdose.

Together, our findings support electrical stunning as a rapid and humane euthanasia method for 4 dpf zebrafish larvae. Electrical stunning did not induce an acute hsp70 stress response or detectable neuronal tissue damage, while largely preserving tissue integrity for post-mortem analyses. The delayed rise in hsp70 suggests that cellular stress responses may continue during and after the process of organismal death. Overall, these results strengthen electrical stunning as a promising alternative to other methods of euthanasia and support further standardization of this technique for broader use.

## Acknowledgements

A.B.A received funding from the German Federal Ministry of Research, Technology and Space under grant number 03LWH0150. Responsibility for the content of this publication lies with the authors.

## Data Availability

All data are available from the authors upon request.

## Author contributions

The conceptualization of the project was done by A.B.A. and D.-S.B. with the help of R.T. and H.-R.K. Experiments were conducted by K.S., M.G. and S.K with the help of D.M. All data were analyzed by D.-S.B. and K.S. Resources were provided by A.B.A, R.T. and H.-R.K. Figures were made by D.-S.B., K.S. and A.B.A. The manuscript was written by D.-S.B. and A.B.A. with the help of R.T. and H.-R.K. The funding acquisition was done by A.B.A. and D.-S.B.

## Competing interests

The authors declare no competing interests.

## Methods

### Histology Experiments

Both *mitfa*^*w2/w2*^ and the transgenic *Tg(elavl3:H2B-GCaMP6f)* line were used in the histology staining experiments. The lines were chosen according to the lines used in the our previous study investigating electrical stunning (Burkhardt et al., 2025). Hematoxylin and eosin staining of sections was applied for both *mitfa*^*w2/w2*^ and transgenic larvae euthanized via electrical stunning as well as for two corresponding control groups which did not undergo any treatment. In total 30 larvae were examined. Electrical stunning was performed using the setup described in Burkhardt et al. (2025) and the proposed parameter set (50 V/cm, SIN, 60Hz, 32 s). Directly after exposure to the electrical field (no exposure for the control groups) larvae were fixed in glutardialdehyde (2 % cacodylate buffered solution) (Eltoum et al., 2001). After fixation, larvae were dehydrated using the following procedure: rinsing in 70 %, 80 %, 90 %, 96 %, 100 % ethanol (3 x 15 min for each concentration). Embedding was done in a synthetic resin (Technovit 7100, Heraeus Kulzer, Germany) overnight before a hardener (Technovit 7100 plus hardener 1, Heraeus Kulzer, Germany) was added in the next morning. For each larva series of 2 µm sections were cut using a rotatory microtome (Leica RM 2265, Wetzlar, Deutschland). Staining of the sections was performed using an automated stainer (Varistain 24-4, Thermo Fisher Scientific, Waltham, USA) and stained slices were covered with a drop of Roti-Hisokitt (Carl Roth GmbH, Karlsruhe, Germany). All slices were examined using an Axioscope 2 microscope (Zeiss, Germany) with 20x and 40x magnification and a camera (AxioCam MRC, Zeiss). All tissue types were examined with regard to disrupted tissue, karyolysis, enlarged intercellular spaces, hypertrophy/atrophy, hemorrhages and degeneration.

**Figure 3.**
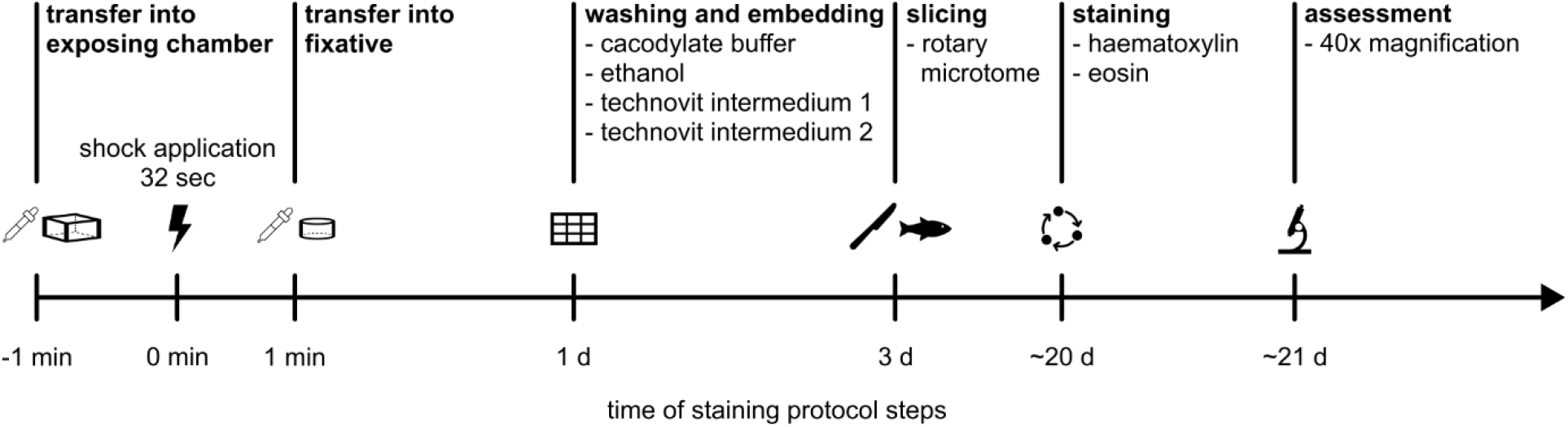
Protocol for the histological examination of electrically stunned zebrafish larvae.

### Hsp70 Experiments

To compare stress responses of zebrafish larvae to different euthanasia methods, induction of hsp70 in *mitfa*^*w2/w2*^ larvae were measured in four treatment groups: anesthetic overdose of MS-222 (1000 g/ml for 60 min), hypothermia (0-2°C for 60min), electrical stunning (AC, SIN, 60 Hz, 50 V/cm for 32 s as proposed in Burkhardt et al., 2025) and heat shock treatment (39°C for 60 min, similar to Murtha and Keller, 2003). In order to compare all the measurements to a baseline, hsp70 measurements were also taken in untreated larvae (Figure 2A). For each group, 200 larvae were used in 10 independent trials with 20 animals per trial. After treatment, larvae were shock frozen in liquid nitrogen and stored at −80°C for hsp70 analysis. Within each trial, larvae were pooled and homogenized in 20 µl extraction buffer according to Scheil et al., 2008. Homogenate was then centrifuged (10 min, 20,000 g at 4°C) and the total protein concentration was determined (Bradford, 1976). From each sample, 20 µg of protein were subjected to SDS-PAGE for 15 min at 80 V and for 1 h at 120 V. The protein was electro-transferred to nitrocellulose for 2h (2mA/cm^2^) and stained by an antibody complex (mouse α human hsp70, peroxidase-conjugated goat α mouse IgG; see Scheil et al., 2008).

Western blot protein bands were first quantified using a densitometric image analysis system (Herolab E.A.S.Y., Germany) and then normed to an internal adult zebrafish standard, run in the parallel on each gel. Outliers with a z-score > 2 were excluded from each group. Significance levels were calculated using Mann-Whitney-Wilcoxon’s U-test as data were not normally distributed (tested by Shapiro-Wilk test).

## Notes

### Competing Interest Statement

The authors have declared no competing interest.

